# EigenBrain maps cortical molecular architecture from functional, geometric, and structural eigenmodes

**DOI:** 10.64898/2026.10.27.758642

**Authors:** Bin Wan, Penghui Du, Asia Ferrari, Amin Saberi, Alexandra John, Jiadong Yan, Xiaobo Liu, Justine Y. Hansen, Zhen-Qi Liu, Seok-Jun Hong, Fabien Carruzzo, Mariia Kaliuzhna, Tal Geffen, Teresa Katthagen, Bratislav Misic, Paul M. Thompson, Dimitri Van De Ville, Daniel S. Margulies, Varun Warrier, Richard A.I. Bethlehem, Florian Schlagenhauf, Stefan Kaiser, Boris C. Bernhardt, Sofie L. Valk, Matthias Kirschner

## Abstract

Neurotransmitter receptor and transporter distributions shape neural communication across the cortex and underpin mechanistic interpretations of brain function. Magnetic resonance imaging (MRI)-derived eigenmodes, low-dimensional spatial patterns that capture the principal axes of cortical organization, are increasingly used to reconstruct brain maps and offer a potential route to approximating molecular topographies in large cohorts where direct measurements are unavailable. Yet whether the type of MRI data used to derive these eigenmodes determines which receptor and transporter distributions can be approximated, and which molecular systems are accessible through this approach at all, remains unknown. Using a spatial reconstruction framework applied to 23 molecular systems spanning neurotransmitter, metabolic, and pathological markers, we show that eight receptor and transporter maps, spanning serotonin, dopamine, histamine, cholinergic, and opioid markers, can be selectively approximated from functional eigenmodes but not structural or geometric ones. This selectivity also reveals that functional brain organization encodes a specific subset of neuromodulatory topographies that structural and geometric organization does not. To illustrate translational potential in clinical cohorts where molecular imaging is unavailable, we applied individual-level eigenmode decompositions of MRI data projected onto the eight validated molecular systems and detected cross-sectional differences in autism and longitudinal changes in schizophrenia. Together, these findings reveal a difference in how geometric, structural, and functional brain organization relate to molecular architecture, offering a method for approximating validated molecular profiles in population neuroimaging cohorts that lack direct molecular imaging.

## Introduction

Neurotransmitter receptor distributions, synaptic density, and metabolic activity jointly shape neural communication, and their disruption relates to a broad range of psychiatric and neurological conditions ^1–18^. For example, excitation-inhibition imbalance alters social and sensory networks in autism ^19–21^; disrupted dopaminergic and glutamatergic signaling converges on prefrontal-limbic circuits in schizophrenia ^22–25^; and in Alzheimer’s disease, amyloid-β and tau spread along connectome pathways in patterns that track cognitive decline ^26–28^. Linking molecular disruptions to their circuit-level consequences in individual patients, rather than in group averages, is therefore a central goal for both biological psychiatry and the development of targeted treatments.

Positron emission tomography (PET) provides the most direct *in vivo* measurements of the brain’s molecular architecture, including receptor densities and pathological protein burden, as well as glucose metabolism as a proxy for regional neuronal activity ^8^. Each scan, however, captures a single tracer and carries high cost and radiation exposure, so multi-marker profiling within one person is rarely feasible ^29,30^, and limited scanner access constrains sample sizes and generalizability ^8^. Deep-learning models can predict some PET-derived maps, notably glucose metabolism, from structural MRI ^31,32^, but they usually target one marker, operate as black boxes, and reveal little about the biological principles that link brain structure to molecular architecture. What is missing is a framework that reconstructs molecular profiles directly from the brain’s intrinsic spatial organization: geometry, white-matter wiring, and functional connectivity. With such an approach, the resulting maps remain biologically interpretable and the reconstruction principle is transparent rather than opaque.

A multiscale and biophysically grounded view of brain organization offers exactly this foundation ^33–38^. At the micro- and mesoscale, distinct neurotransmitter systems define the molecular identity of local circuits (**Fig. 1a–b**), circuits whose energetic demand is tracked by cerebral glucose metabolism ^39^, and whose signaling is impaired when the microenvironment is disrupted by pathological processes including neuroinflammation, synaptic loss, protein aggregation, and myelin degeneration (**Fig. 1c)**. At the macroscale, these substrates give rise to coordinated activity measurable as functional connectivity (FC), constrained by cortical geometry (geodesic distance, GD) and white-matter wiring (structural connectivity, SC) (**Fig. 1d**). A single framework connecting molecular and macroscale brain organization must therefore span biophysical scales while remaining biologically interpretable and tractable enough to apply across individuals and datasets ^36,40^.

**Fig. 1:**
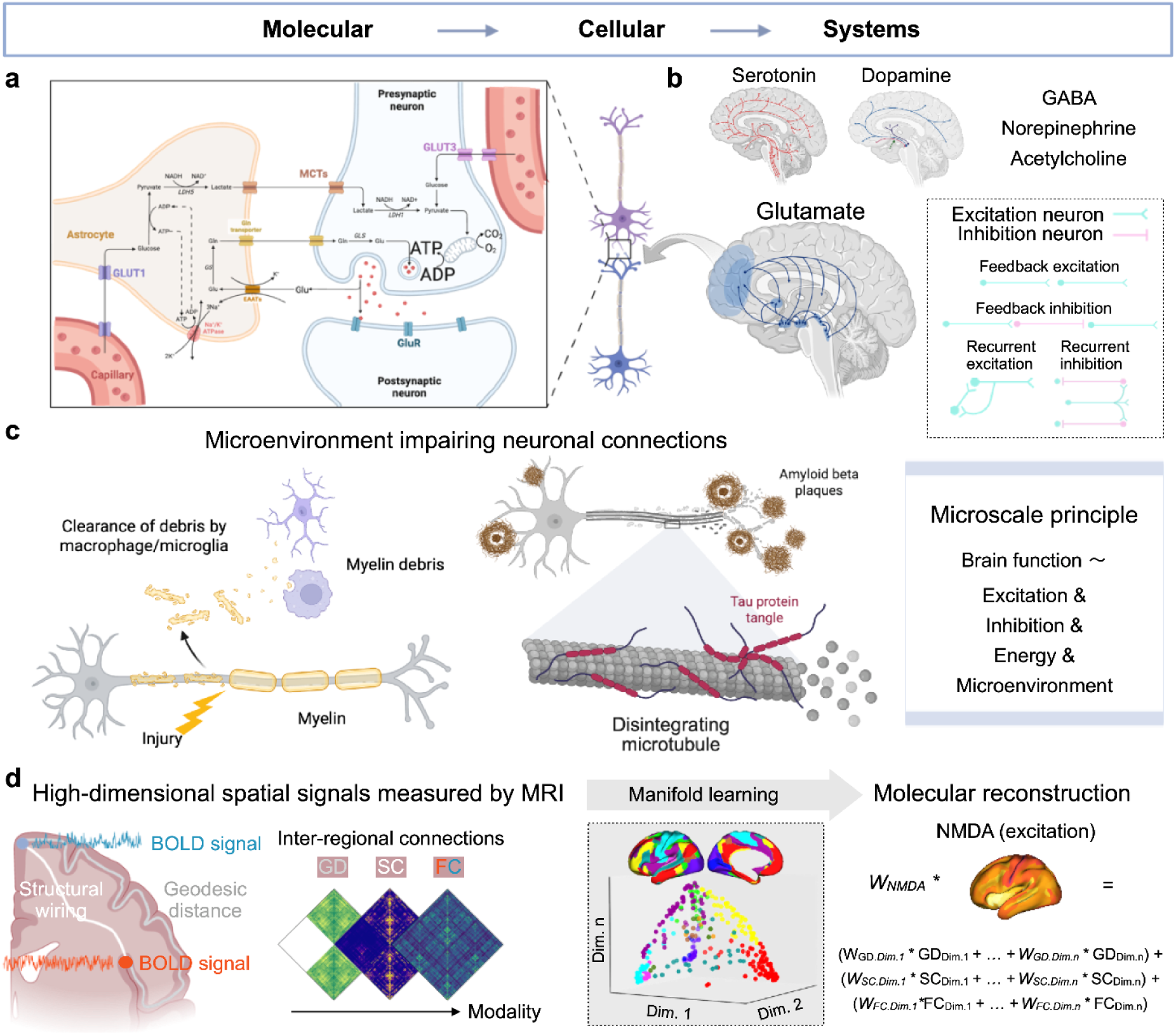
Conceptual overview of EigenBrain. **a,** The glutamate-glutamine cycle at an excitatory synapse, illustrating the energetic demands of neurotransmitter synthesis, release, and recycling between neurons and astrocytes. **b,** Glutamatergic projections form the brain’s principal excitatory system, alongside neuromodulatory systems including serotonergic, dopaminergic, GABAergic, noradrenergic, and cholinergic pathways that regulate local circuit function through excitation-inhibition balance. **c,** Representative pathological disruptions of synaptic transmission, including demyelination following axonal injury, amyloid-β plaque accumulation, and tau tangle formation, each impairing neuronal signaling through distinct mechanisms. The right panel summarizes the microscale principle that brain function depends on excitation, inhibition, energy supply, and microenvironmental integrity. **d,** Multimodal MRI captures complementary spatial signals: cortical geodesic distance (GD), structural connectivity (SC) from white-matter tractography, and functional connectivity (FC) estimated from time-series correlation of blood-oxygen-level-dependent (BOLD) signals across regions. These define GD, SC, and FC matrices, from which manifold learning extracts low-dimensional eigenvectors representing the principal axes of brain spatial organization. EigenBrain reconstructs whole-brain molecular maps as weighted linear combinations of these eigenvectors, illustrated here for the N-methyl-D-aspartate (NMDA) receptor.

Manifold learning, a dimensionality reduction approach widely applied across genomics, single-cell biology, and systems neuroscience to extract low-dimensional structure from high-dimensional data, offers a principled route to this goal ^41^. These patterns, termed gradients or eigenmodes, form a compact basis for reconstructing complex brain maps as weighted combinations of a small number of modes. Geometric eigenmodes reconstruct task-evoked fMRI activation ^42^, and functional gradients account for hierarchical variation in cerebral energy metabolism ^43^. Together these results suggest that MRI-derived eigenmodes could provide a general spatial basis for reconstructing molecular profiles across the cortex, turning individual-level molecular mapping into an inference problem on standard MRI rather than a PET acquisition problem. Whether structural or functional eigenmodes are better suited to this task, however, and which molecular systems are recoverable, remains incompletely understood.

Identifying which MRI modality relates to molecular architecture addresses only part of the problem; a second concern is how molecular information is currently applied in clinical neuroscience. The dominant approach correlates group-level cortical abnormality patterns against group-level normative neurotransmitter atlases to identify candidate molecular substrates ^4,9,44–52^. This strategy has successfully associated disease-related cortical alterations to different receptor systems, but only at a group level. Both the abnormality map and the molecular reference are population averages, so the substrates identified describe a canonical patient rather than any individual. Resolving modality selection and extending molecular reconstruction from group- to individual-level are therefore necessary conditions for translating molecular mapping into individual-level inferences which could be clinically useful.

To address these questions, we developed EigenBrain, a spatial reconstruction framework that reconstructs molecular maps from normative MRI connectome eigenmodes at the group-level, then transfers these reconstruction weights to individual MRI data to generate subject-specific whole-brain molecular profiles without requiring PET imaging (**Fig. 1**). We applied eigenmodes derived from three spatial bases, including GD, SC, and FC, to reconstruct cortical molecular profiles from multimodal MRI through EigenBrain. We built reconstructions against 20 neurotransmitter receptor and transporter maps from Neuromaps ^34^ using group-level Human Connectome Project (HCP) data ^53,54^, then extended to individual-level validation in two OpenNeuro datasets with paired PET and MRI: glucose metabolism from the energy cost dataset ^55^, and tau and amyloid-β from the Dallas Lifespan Brain Study ^56^. This design allowed us to ask, first, which modality yields eigenmodes that capture molecular architecture more accurately; second, which molecular systems carry genuine signal in the MRI-derived eigenmodes rather than simply reflecting the smooth spatial structure of the cortex; and third, whether eigenmode-based molecular profiles capture clinically relevant individual variability. Answers to these questions establish the degree to which functional brain organization relates to molecular architecture and whether that encoding is reliable enough to serve as a surrogate for direct molecular imaging in large-scale MRI cohorts.

## Results

### Reconstructing 20 neurotransmitter receptor and transporter maps

We first tested whether EigenBrain could reconstruct group-level neurotransmitter receptor and transporter distributions from multimodal MRI, using 20 molecular maps from Neuromaps and MRI from unrelated HCP participants (*n* = 94; **Fig. 2a**). From T1-weighted MRI, diffusion-weighted imaging, and resting-state fMRI, we derived GD, SC (streamline counts), and cortical FC, respectively (**Fig. 2b**). To reduce computational cost, we summarized all data with a multimodal parcellation of 180 cortical regions per hemisphere ^57^. For each modality, diffusion map embedding ^58,59^, implemented in BrainSpace ^41^, extracted low-dimensional eigenvectors representing the gradients of brain spatial organization (**Fig. 2c**). Based on prior evidence that reconstruction convergence (defined as the elbow point) occurs at approximately 70 components ^43^, we extracted the first 100 eigenvectors from each modality to ensure adequate coverage across all molecular maps.

**Fig. 2:**
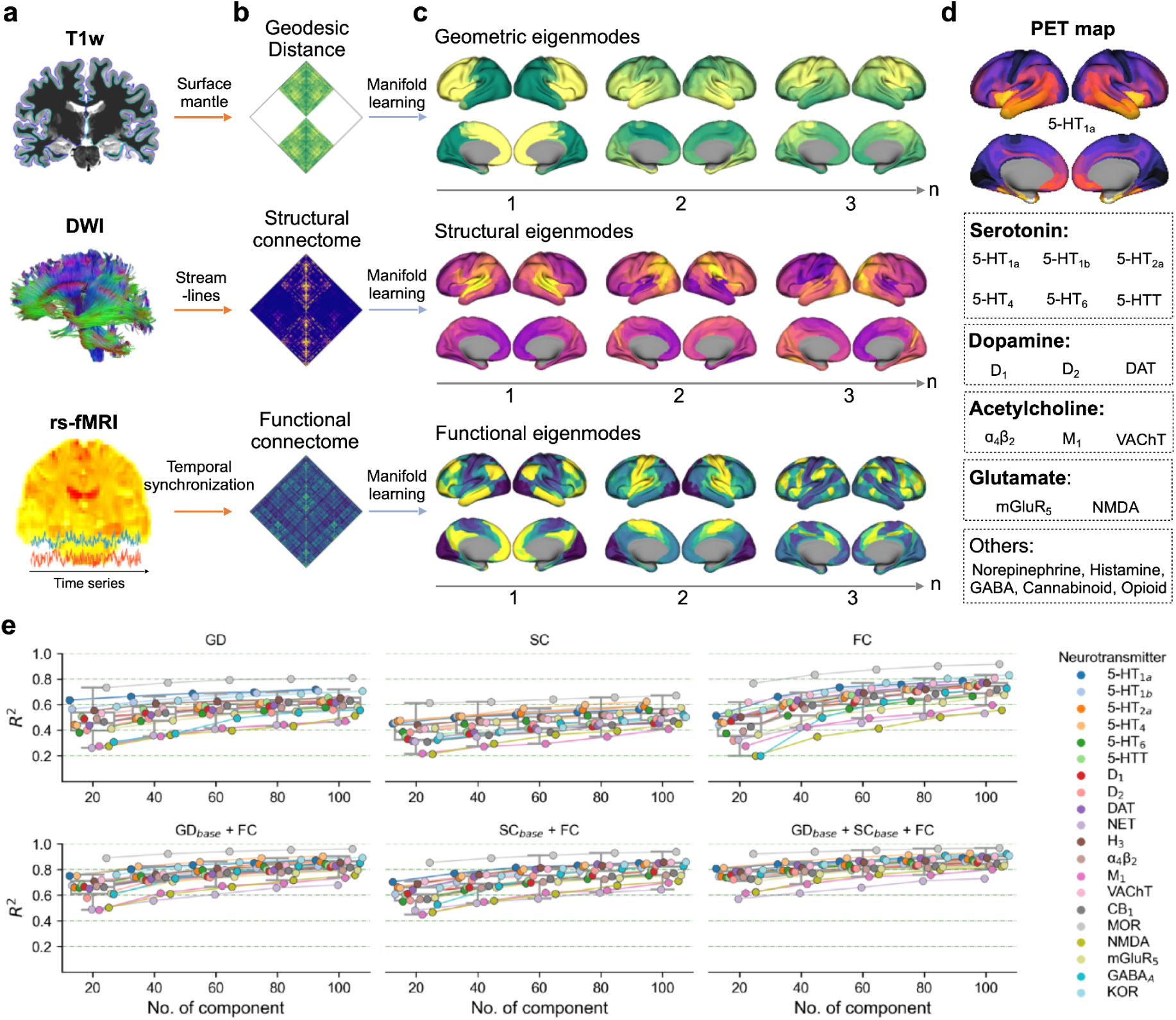
Reconstructing 20 neurotransmitter receptor and transporter maps. **a**, T1-weighted MRI, diffusion-weighted imaging, and resting-state fMRI were used to derive geodesic distance (GD), white-matter structural connectivity (SC), and cortical functional connectivity (FC), respectively (**b**). **c**, Diffusion map embedding was applied to each matrix to extract low-dimensional eigenvectors, shown here as the first three geometric, structural, and functional gradients. **d,** Group-level PET- and SPECT--derived neurotransmitter receptor and transporter maps from Neuromaps, illustrated for 5-HT_1a_, were reconstructed with EigenBrain models based on GD, SC, FC, GD_base_ + FC, SC_base_ + FC, and GD_base_ + SC_base_ + FC. The 20 maps span serotonergic, dopaminergic, cholinergic, glutamatergic, noradrenergic, histaminergic, GABAergic, cannabinoid, and opioid systems. **e,** Reconstruction performance, quantified by *R*², as a function of the number of eigenvector components in each model. FC provided the strongest single-modality reconstruction, while multimodal models combining FC with GD_base_ and/or SC_base_ further improved accuracy.

The 20 molecular maps (**Fig. 2d**) comprised six serotonergic (5-HT_1a_, 5-HT_1b_, 5-HT_2a_, 5-HT_4_, 5-HT_6_, and 5-HTT), three dopaminergic (D_1_, D_2_, and DAT), three cholinergic (ɑ_4_β_2_, M_1_, and VAChT), two glutamatergic (mGluR_5_ and NMDA), one noradrenergic (NET), one histaminergic (H_3_), one GABAergic (GABA_A_), one cannabinoid (CB_1_), and two opioid maps (MOR and KOR). We fitted six reconstruction models: three single-modality models (GD, SC, or FC eigenvectors), and three multimodal models combining FC with GD_base_, SC_base_, or both. Here “base” denotes the modality-specific component count at the elbow of the component-number/*R²* curve (**Fig. 2e** and **Source Data**).

Across the 20 molecular maps, SC converged with the fewest components (31 ± 13 on average) but achieved lower reconstruction accuracy (Pearson *r* = 0.65 ± 0.06) than GD and FC. GD required 39 ± 13 components and reached higher accuracy (Pearson *r* = 0.73 ± 0.07), whereas FC required 42 ± 14 components and achieved the highest single-modality accuracy (*r* = 0.79 ± 0.07). The number of components needed for convergence was significantly lower for SC than for GD (*t* = 2.263, *P* = 0.034) or FC (*t* = 3.494, *P* = 0.002). At a fixed dimensionality of 40 components, FC again performed best, with mean *R*² = 0.58 ± 0.10 versus GD (*R*² = 0.52 ± 0.11; FC versus GD: *t* = 5.308, *P* < 0.001) and SC (*R*² = 0.43 ± 0.08; FC versus SC: *t* = 8.720, *P* < 0.001). FC thus provided the most accurate single-modality basis, while GD and SC captured complementary spatial structure. In addition, we also observed that the reconstruction was not significantly related to the sample size (Spearman *r*: 0.12 to 0.19 for each modality with all *P* values > 0.4).

To test whether structural and geometric information contributes to molecular reconstruction beyond functional organization alone, we used GD and SC eigenmodes as a fixed spatial base and asked whether adding FC eigenmodes improved accuracy. GD_base_ combined with FC converged at 55 ± 24 FC components and achieved accuracy of 0.87 ± 0.05; SC_base_ with FC converged at 53 ± 16 FC components and achieved accuracy of 0.86 ± 0.05; and GD_base_, SC_base_, and FC together required 55 ± 23 FC components and achieved the highest overall accuracy (*r* = 0.90 ± 0.04). The three multimodal models did not differ significantly from one another (all *P* > 0.05), indicating that the complementary variance contributed by GD and SC reaches a ceiling regardless of whether one or both of the geometric and structural modalities are included. This selectivity suggests that GD and SC capture largely overlapping spatial variance beyond FC. These results indicate that FC provides the strongest single-modality basis, that structural and geometric organization contribute complementary but partially redundant spatial information, and that the full multimodal model offers the most complete reconstruction of cortical molecular topography, with incremental but significant gains over FC alone (reconstruction maps in **Supplementary Fig. S1-20**).

Next, it is essential to identify which molecular reconstructions reflect connectome-specific information rather than inflation by spatial autocorrelation. For each molecular marker, we generated 1,000 surrogate maps using spatial variogram models that preserve the empirical spatial autocorrelation of the true map ^60^. Each surrogate was reconstructed using the same EigenBrain procedure, and the accuracy of the true reconstruction was compared with the resulting null distribution. The analyses were performed for single-modality GD, SC, and FC models (**Supplementary Fig. S21-40**). Across the 20 maps, FC reconstructions exceeded the spatially constrained variogram null for a subset of systems, whereas GD and SC did not. In particular, all single-modality GD and SC reconstructions fell within their surrogate distributions (all *P*_variogram_ > 0.05), indicating that at the single-modality group level their accuracy was not separable from spatial smoothness. By contrast, FC reconstruction exceeded the surrogate null for 8 of the 20 maps: 5-HT_1a_ (components ≥ 40), 5-HTT (≥ 100 components), D_2_ (≥ 100 components), DAT (≥ 60 components), H_3_ (≥ 100 components), VAChT (≥ 20 components), MOR (≥ 20 components), and KOR (≥ 40 components).

### Reconstructing glucose metabolism, amyloid-β, and tau at both group and individual levels

We next extended EigenBrain beyond receptor and transporter maps to three PET-derived metabolic and pathological profiles for which group-level and individual-level data were available: glucose metabolism, amyloid-β deposition, and tau deposition. For glucose metabolism, we used individual-level FDG-PET and multimodal MRI from the Munich dataset (*n* = 19) ^55^ and computed the cerebral metabolic rate of glucose (CMRglc; **Fig. 3a**). For amyloid-β and tau, we used the third wave of the Dallas Lifespan Brain Study (*n* = 61) (Park et al., 2025) comprising [18F]-AV45 PET, [18F]-AV1451 PET, and multimodal MRI, and computed standardized uptake value ratio (SUVR) maps (**Fig. 3b**). In the Dallas sample, cognitive impairment (Alzheimer’s Disease Assessment Scale, ADAS, total score) correlated with age (Pearson *r* = 0.53, *P* < 0.001, **Fig. 3b**).

**Fig. 3:**
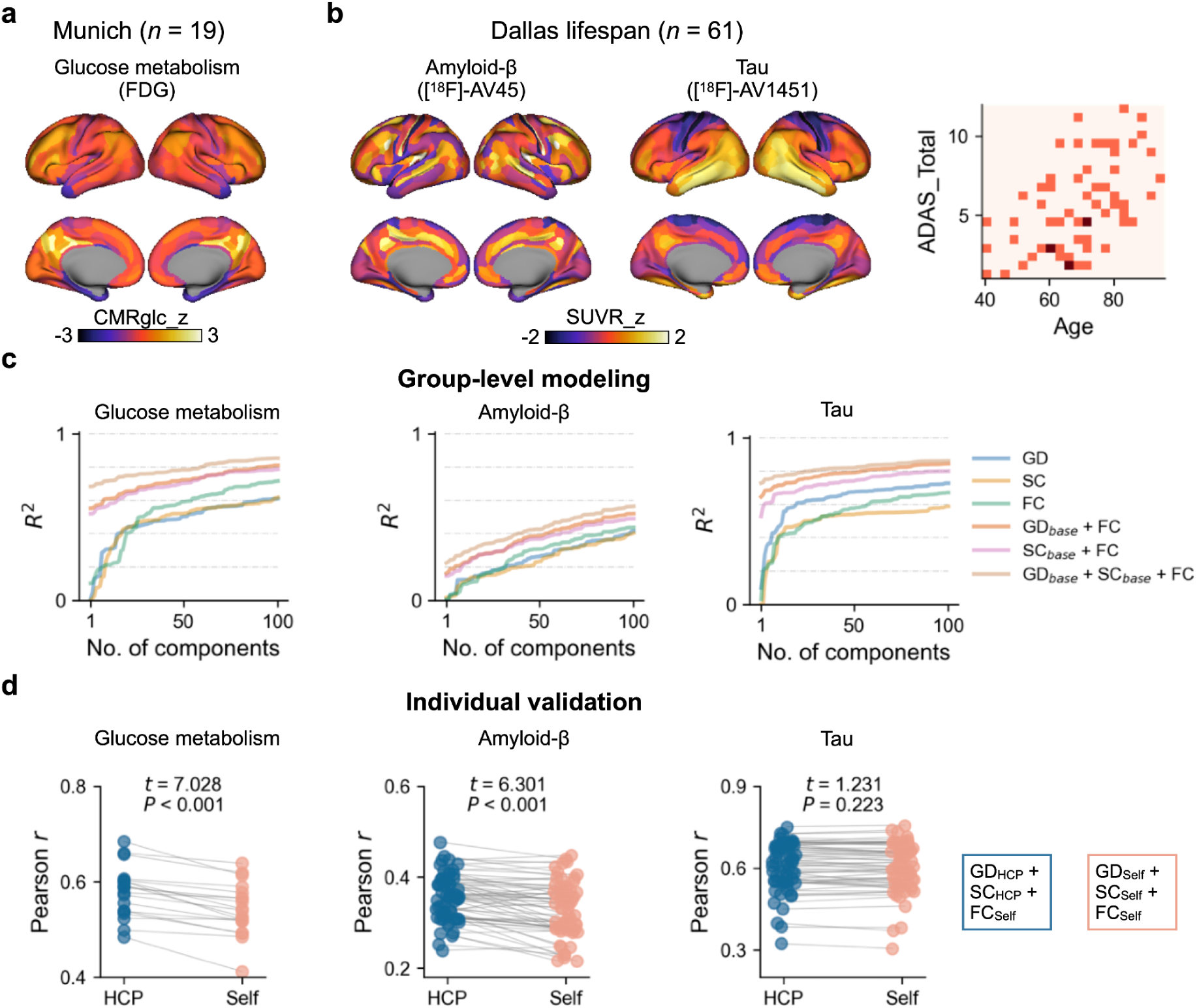
Reconstructing glucose metabolism, amyloid-β deposition, and tau deposition. **a,** Group-average cerebral metabolic rate of glucose (CMRglc) map derived from FDG-PET in the Munich dataset (*n* = 19). **b,** Group-average amyloid-β and tau deposition maps derived from 18F-AV45 and 18F-AV1451 PET, respectively, in wave 3 of the Dallas Lifespan Brain Study (*n* = 61). **c,** Group-level reconstruction performance for glucose metabolism, amyloid-β, and tau as a function of the number of eigenvector components included in the model. Models included GD, SC, FC, GD_base_ + FC, SC_base_ + FC, and GD_base_ + SC_base_ + FC. Multimodal models showed the highest performance for glucose metabolism and tau, whereas amyloid-β showed weaker reconstruction and did not reach clear convergence within 100 components. **d,** Individual-level validation comparing reconstruction accuracy between models using HCP-derived GD and SC with participant-specific FC (GD_HCP_ + SC_HCP_ + FC_Self_) and models using participant-specific GD, SC, and FC (GD_Self_ + SC_Self_ + FC_Self_). Accuracy was quantified by Pearson correlation between reconstructed and true PET-derived maps. Abbreviations: ADAS, Alzheimer’s Disease Assessment Scale; CMRglc, cerebral metabolic rate of glucose; FC, functional connectivity; FDG, fluorodeoxyglucose; GD, geodesic distance; SC, structural connectivity; SUVR, standardized uptake value ratio.

We first trained group-level models from the mean PET map and group-level HCP eigenvectors (**Fig. 3c**), then applied the learned weights to individual participants. For CMRglc, GD converged at 21 components (*R*^2^ = 0.42), SC at 26 (*R*^2^ = 0.46), and FC at 59 (*R*^2^ = 0.62). Multimodal models improved performance: GD_base_ + FC required 77 FC components (*R*^2^ = 0.78), SC_base_ + FC at 74 (*R*^2^ = 0.77), and GD_base_ + SC_base_ + FC at 74, reaching the highest performance (*R*^2^ = 0.83). Amyloid-β reconstruction was weaker, and no model reached a clear convergence point within 100 components (*R*^2^_Max_ = 0.43 for single modality and *R*^2^_Max_ = 0.56 for combined modality), consistent with its more diffuse, spatially heterogeneous distribution. For tau, GD converged at 25 components (*R*^2^ = 0.62), SC at 20 (*R*^2^ = 0.48), and FC at 59 (*R*^2^ = 0.61); multimodal models again improved performance: GD_base_ + SC_base_ + FC was highest at 37 FC components (*R*^2^ = 0.83).

We then assessed individual-level accuracy using the converging number of components from the group-level. Because open datasets do not always provide GD or SC matrices, we compared two variants (**Fig. 3d**): HCP-derived GD and SC combined with participant-specific FC (GD_HCP_ + SC_HCP_ + FC_Self_), and fully participant-specific eigenvectors (GD_Self_ + SC_Self_ + FC_Self_). For glucose metabolism, individual accuracy (Pearson *r* between true and predicted maps) ranged from 0.481 to 0.677 (self-based) and from 0.484 to 0.683 (HCP-based), with the HCP-based model performing significantly better (*t* = 7.028, *P* < 0.001). For amyloid-β, accuracy ranged from 0.230 to 0.448 (self-based) and 0.239 to 0.477 (HCP-based), again favoring the HCP-based model (*t* = 6.301, *P* < 0.001). The lower bound of this range approaches chance-level reconstruction for some individuals, consistent with the absence of a clear convergence point in the group-level model and reflecting the spatially diffuse and heterogeneous distribution of amyloid-β deposition, which lacks the low-dimensional cortical structure that eigenmode-based reconstruction requires. For tau, accuracy ranged from 0.370 to 0.738 (HCP-based) and 0.384 to 0.726 (self-based), with no significant difference (*t* = 1.231, *P* = 0.223). Together, EigenBrain reconstructs structured glucose-metabolism and tau maps from MRI-derived eigenvectors provided that individual FC matrices are available. Reconstruction was weaker for amyloid-β, likely reflecting its diffuse spatial distribution. It suggests that diffuse signals such as amyloid-β may require more flexible individual-level, deep-learning, and higher resolution MRI input approaches.

At the group level, spatial autocorrelation testing using HCP-derived eigenmodes revealed no surviving maps among the metabolic and pathological markers (**Supplementary Fig. S41-43**), indicating that group-level functional eigenmodes do not capture glucose metabolism, amyloid-β, or tau beyond cortical smoothness. This null result does not preclude recoverability under different conditions: for glucose metabolism specifically, our previous work demonstrated that reconstruction from sample-specific FC eigenvectors, derived from the same individuals with paired PET data, does exceed the spatial autocorrelation null ^43^. These findings suggest that individual-level eigenmodes carry molecular information that group-level templates do not.

Together, these results show that EigenBrain reconstruction is molecular-system specific. At the group-template level, FC eigenvectors capture the cortical topography of a defined subset of neurotransmitter systems beyond spatial autocorrelation, whereas GD and SC eigenvectors do not. Because the variogram test is applied to fixed group-level templates, it establishes group-level spatial specificity rather than direct individual molecular specificity. These findings therefore define the appropriate scope of future EigenBrain applications: for MRI-only cohorts that rely on HCP group-level eigenvectors for molecular translation, analyses should be restricted to the eight neurotransmitter systems validated beyond the spatial nulls. In datasets with paired subject-level PET or other molecular imaging, reconstruction should instead be tested directly within that dataset using sample-specific eigenvectors and the same spatial null framework, because additional systems may be recoverable in a sample-specific setting.

### FC-informed molecular profiles in autism and schizophrenia

Next, to illustrate the translational potential of EigenBrain in clinical settings, we examined two psychiatric conditions with well-characterized cortical abnormalities and hypothesized neuromodulatory involvement: autism spectrum disorder and schizophrenia. These disorders represent complementary use cases, with autism enabling cross-sectional case-control inference across a large multi-site sample and schizophrenia enabling longitudinal tracking of molecular profiles within individuals over time. So the reconstructed profiles could not be validated against direct receptor measurements, and the findings should be interpreted as hypothesis-generating rather than as evidence of true molecular alteration. Using trained reconstruction weights from HCP and subject-level FC matrices, we generated the eight predicted molecular signaling maps for each participant (**Fig. 4a**).

**Fig. 4:**
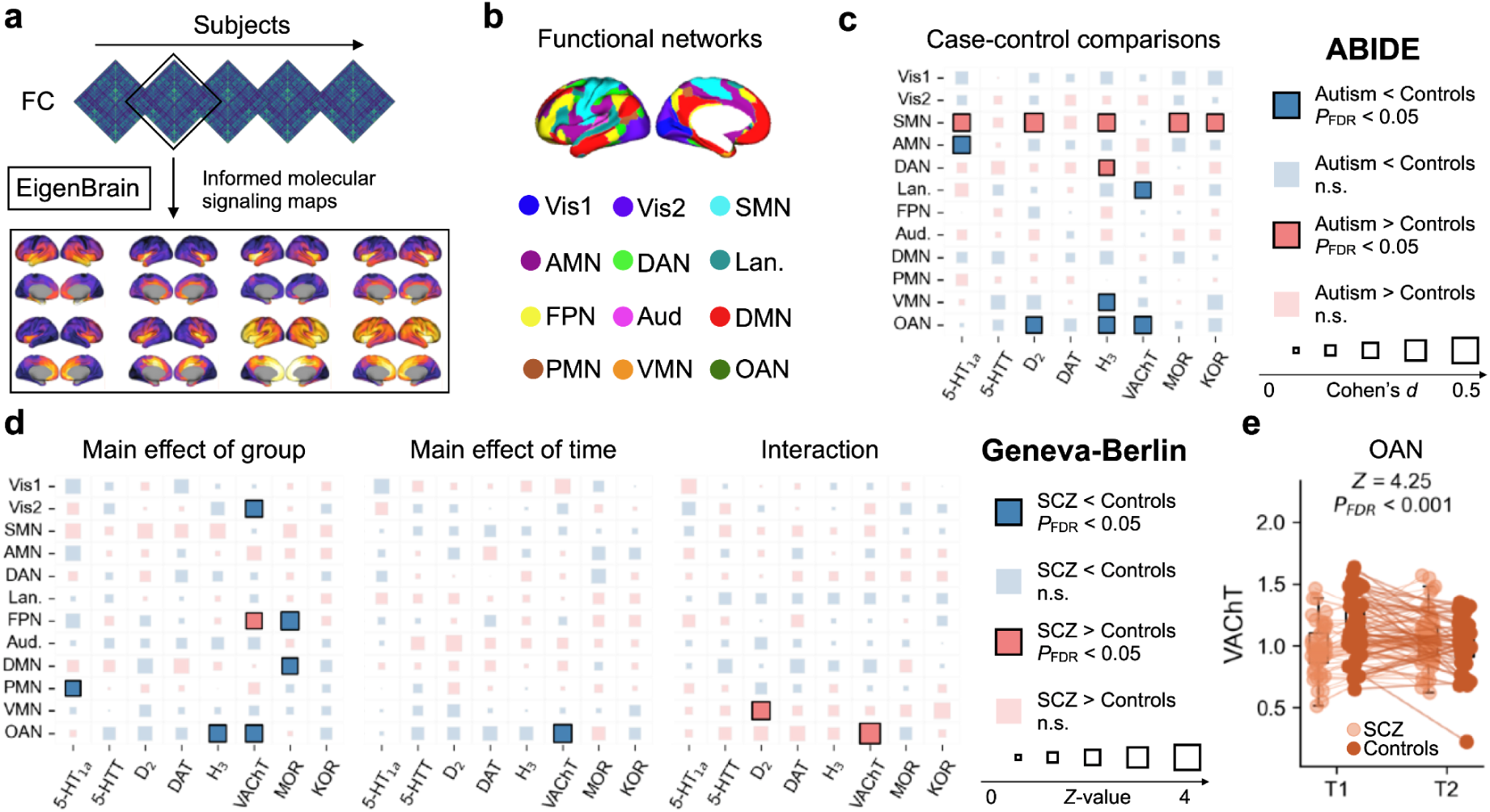
Informed molecular signaling in autism and schizophrenia. **a,** Subject-level FC matrices were entered into the EigenBrain framework to derive the eight validated FC-informed molecular maps for each participant. **b,** Parcel-wise informed molecular signaling values were summarized across 12 functional networks including primary visual (Vis1), secondary visual (Vis2), somatomotor (SMN), action-mode (AMN), dorsal attention (DAN), language (Lan.), frontoparietal (FPN), auditory network (Aud), default mode (DMN), posterior multimodal (PMN), ventral multimodal (VMN), orbito-affective (OAN). **c,** Network-level case-control differences between autism and controls; square color indicates direction and significance, square size indicates effect size. **d,** Network-level differences between individuals with schizophrenia and controls using generalized estimating equations in the Geneva-Berlin longitudinal cohort. **e,** Significant interaction was found in the OAN, showing different longitudinal change between schizophrenia and controls.

For autism, we analyzed five Autism Brain Imaging Data Exchange (ABIDE) sites ^61,62^, yielding 140 autistic individuals and 143 non-autistic controls, and reconstructed individual neurotransmitter profiles from HCP-derived GD and SC eigenvectors with participant-specific FC. Across the 8 maps, case-control differences were found for D_2_ in the cingulate cortex (**Supplementary Fig. S44**). We then summarized predicted molecular signaling values into 12 functional networks (**Fig. 4b**) defined by ^63^: primary visual (Vis1), secondary visual (Vis2), somatomotor (SMN), cingulo-opercular/action-mode (AMN) ^64^, dorsal attention (DAN), language (LAN), frontoparietal (FPN), auditory network (AUD), default mode (DMN), posterior multimodal (PMN), ventral multimodal (VMN), orbito-affective (OAN). Network-level analyses revealed a concentrated pattern of increased molecular signaling in SMN in autism, involving 5-HT_1a_ (*d* = 0.35, *P*_FDR_ = 0.019), D_2_ (*d* = 0.42, *P*_FDR_ = 0.005), H_3_ (*d* = 0.36, *P*_FDR_ = 0.034), MOR (*d* = 0.43, *P*_FDR_ = 0.005), and KOR (*d* = 0.38, *P*_FDR_ = 0.020), and decreased signaling in OAN in autism, involving D_2_, H_3_, and VAChT (**Fig. 4c**; statistics in **Source Data**). To verify that the SMN molecular differences were not driven by SMN FC (**Supplementary Fig. S45**), we controlled for mean SMN-FC in our model and observed similar molecular effects with significant D_2_ (*d* = 0.37, *P*_FDR_ = 0.022) and MOR (*d* = 0.39, *P*_FDR_ = 0.015), and trend significance of 5-HT_1a_ (*d* = 0.31, *P*_FDR_ = 0.052), H_3_ (*d* = 0.33, *P*_FDR_ = 0.066), and KOR (*d* = 0.34, *P*_FDR_ = 0.055). This suggests that the molecular reconstruction carries information beyond general FC spatial structure, rather than simply reflecting broad connectivity patterns.

For schizophrenia, the Geneva-Berlin longitudinal cohort comprised 47 individuals with schizophrenia and 45 controls assessed at two time points with repeat assessments approximately three months apart. Using generalized estimating equations at the network level, the main effect of group showed decreased 5-HT_1a_ in PMN (*Z* = -2.41, *P*_FDR_ = 0.047), decreased H_3_ in OAN (*Z* = -2.93, *P*_FDR_ = 0.041), decreased VAChT in Vis2 (*Z* = -3.06, *P*_FDR_ = 0.023) and OAN (*Z* = -3.00, *P*_FDR_ = 0.016), increased VAChT in FPN (*Z* = 2.73, *P*_FDR_ = 0.025), and decreased MOR in FPN (*Z* = -3.28, *P*_FDR_ = 0.012) and DMN (*Z* = -2.83, *P*_FDR_ = 0.027), in schizophrenia (**Fig. 4d**). Time-varying effects appeared for VAChT in OAN. The significant interaction involved VAChT (*Z* = 4.25, *P*_FDR_ < 0.001) in OAN (**Fig. 4e**) and D_2_ in VMN (*Z* = 3.24, *P*_FDR_ = 0.014). These findings remained robust when controlling for their network FC except for the VAChT in FPN (**Supplementary Fig. 45**). All the statistics can be seen in **Source Data**. These profiles suggest both persistent and dynamic components in schizophrenia, although larger samples and longer follow-up are needed to separate stable pathology from stage-dependent change.

## Discussion

Through systematic alignment of connectome eigenmodes from multimodal MRI with PET- and SPECT-derived molecular distribution maps across 23 systems, we show that functional eigenmodes outperform structural and geometric eigenmodes as a spatial basis for molecular reconstruction, and that participant-specific GD and SC eigenmodes offer no advantage over normative HCP-derived templates. Of the 23 systems tested, eight neuromodulatory topographies were reconstructed beyond spatial autocorrelation using functional eigenmodes, identifying a validated subset of molecular systems as candidates for individual-level reconstruction in large-scale MRI cohorts. Extending to metabolic and pathological markers, individual-level reconstruction of glucose metabolism and tau from participant-specific FC eigenmodes achieved accuracy comparable to group-level performance, while amyloid-β, whose deposition is spatially diffuse, resisted reconstruction at any component number, demonstrating that individual reconstructability depends on the spatial structure of the molecular map rather than its biological category. As a proof of concept, subject-level FC-informed molecular profiles detected cross-sectional differences in autism and longitudinal changes in schizophrenia, illustrating how molecularly specific hypotheses can be generated in clinical cohorts without direct molecular imaging. Together, these findings reveal a fundamental asymmetry between functional and structural brain organization in their relationship to molecular architecture: the cortical distribution of specific neuromodulatory markers is preferentially encoded in large-scale patterns of spontaneous neural activity, and this encoding is specific enough to serve as a principled basis for imputing validated molecular topographies into large MRI cohorts that lack PET.

Why functional eigenmodes outperform structural ones for molecular reconstruction remains an open question, though several mechanistic interpretations are plausible. Functional connectivity reflects the net output of local neurochemical environments including synaptic receptor densities, neuromodulatory tone, and excitation-inhibition balance, integrated over time and across spatial scales ^8,14,19,21,40,43,55,59^. Structural connectivity and cortical geometry constrain but do not uniquely determine the molecular milieu: although geometric eigenmodes account for large-scale functional activity patterns ^42^, they do not capture the molecular specificity that functional eigenmodes encode, suggesting that spontaneous neural dynamics carry molecular information beyond what anatomy dictates. The fact that FC eigenmodes outperform GD and SC for a specific subset of systems, while multimodal models incorporating GD and SC further improve overall reconstruction accuracy, is consistent with a hierarchical relationship in which structural organization defines a coarse scaffold within which functional dynamics encode finer-grained molecular specificity. This interpretation is supported by evidence that neurotransmitter receptor distributions are shaped by both laminar cytoarchitecture ^8,35,36^ and long-range neuromodulatory projections that drive correlated activity across structurally distant regions ^65^. However, this interpretation warrants caution on two grounds. First, the superiority of FC eigenmodes may partly reflect a methodological advantage: FC matrices are derived from rich temporal data across all region pairs and carry more spatial degrees of freedom than GD or SC, which could inflate reconstruction performance independently of any biological correspondence. Second, current diffusion tractography methods are known to underestimate long-range structural connections, and are prone to false-negative and false-positive reconstructions, particularly in regions with crossing fibers. It means that SC eigenmodes may be limited by measurement error rather than a true absence of structural encoding of molecular architecture. Whether more accurate SC estimation, from higher angular resolution diffusion imaging or next-generation tractography algorithms, would narrow the performance gap with FC remains an important open question.

The eight maps reconstructed beyond spatial autocorrelation span serotonergic (5-HT_1a_, 5-HTT), dopaminergic (D_2_, DAT), histaminergic (H_3_), cholinergic (VAChT), and opioid (MOR, KOR) receptors and transporters, yet other maps within the same families were not, including four additional serotonergic subtypes, D_1_, NET, and two cholinergic markers, indicating that successful reconstruction reflects properties of the specific receptor or transporter map rather than the neurotransmitter family it belongs to. The non-reconstructable maps included both ionotropic receptors such as NMDA and GABA_A_ and metabotropic receptors such as mGluR_5_ and CB_1_ ^8^, indicating that neither a fast versus slow signaling distinction nor an ionotropic versus neuromodulatory classification separates the two groups accurately. What distinguishes reconstructable from non-reconstructable maps remains an open question that systematic analysis across a broader range of molecular markers and normative PET samples will be needed to resolve. Map-level properties including spatial signal-to-noise, spatial autocorrelation structure, and participant age at acquisition remain candidate explanations, though we found no significant relationship between reconstruction success and normative PET sample size or alignment with the FC G1, suggesting that neither data quantity nor gross gradient correspondence determines recoverability. Disentangling the remaining factors will require systematic analysis across a broader range of molecular maps and larger normative PET samples.

The extension to glucose metabolism, amyloid-β, and tau deposition reveals that reconstructability depends not on the biological nature of the marker but on the spatial structure of its cortical distribution. Glucose metabolism and tau, despite belonging to entirely different biological categories, share a spatially structured cortical topography that is well-captured by a low-dimensional eigenmode basis: one is a marker of regional energetic demand ^39,55^, the other a pathological protein whose deposition follows connectome-constrained spreading patterns ^27,28^. Both were reconstructable with individual-level accuracy using HCP-derived templates. Amyloid-β, whose deposition is more spatially diffuse and heterogeneous, resisted reconstruction at any component number. The relevant criterion for eigenmode-based prediction is therefore spatial organization rather than biological category, and reconstructability should be evaluated empirically for each marker rather than assumed from its molecular identity. Deep learning models, which have been successful in predicting FDG-PET from structural MRI ^31,32^, may offer a complementary route for spatially irregular signals, trading the biological interpretability of the eigenmode framework for greater flexibility. A productive synthesis may therefore combine both approaches: eigenmode reconstruction as a biologically grounded baseline for spatially structured systems, and residual or nonlinear prediction for the variance that linear modes cannot explain.

A related approach, REACT (Receptor-Enriched Analysis of functional Connectivity by Targets), uses group-level PET receptor maps as spatial weighting functions to extract receptor-enriched BOLD time series and compute receptor-specific functional connectivity networks for individual participants ^66^. REACT and EigenBrain share the motivation of bringing molecular specificity into individual fMRI analyses, but their directionality is opposite: REACT takes PET maps as input and produces molecularly weighted FC as output, while EigenBrain takes FC eigenmodes as input and reconstructs molecular topographies as output, without requiring any molecular imaging at inference time. A further distinction is that EigenBrain explicitly tests whether each reconstructed map exceeds a spatial autocorrelation null, establishing which systems are recoverable with confidence and which are not. The two approaches are therefore complementary: REACT characterizes receptor-specific connectivity patterns when a normative PET atlas is available, while EigenBrain is designed for cohorts where molecular imaging is absent.

The proof-of-concept analyses in autism and schizophrenia illustrate how FC-informed molecular profiles can generate molecularly specific hypotheses in clinical cohorts without PET. In autism, the concentrated pattern of increased molecular signaling in the somatomotor network, spanning serotonin, dopamine, histamine, and opioid systems, suggests a convergent alteration in the neuromodulatory milieu of somatomotor cortex. Direct *in vivo* PET characterization of cortical receptor systems in autism is limited, largely because existing studies have focused on subcortical regions using tracers with low cortical signal-to-noise, and hypothesis-driven ROI designs would not have detected the cortex-wide somatomotor pattern observed here ^67–69^. The MOR finding is partially corroborated by a recent whole-brain PET study reporting cuneal and precuneal upregulation of MOR availability in autism ^67^, a region that overlaps anatomically with the somatomotor network. Decreased D_2_ signaling in the orbito-affective network is broadly consistent with reduced striatal D_2_ availability reported in the same study, extending that finding to a cortical attentional-affective circuit. For 5-HT_1a_, H_3_, KOR, and VAChT, direct *in vivo* human PET evidence in autism is absent, and the somatomotor alterations identified here remain without a comparator, a gap that underscores both the novelty of the finding and the need for cortex-wide PET studies. In schizophrenia, the longitudinal pattern of decreased serotonergic and opioid signaling in transmodal networks alongside time-varying cholinergic changes in the orbito-affective network is consistent with multi-system neuromodulatory disruption ^12,22,25^, and the interaction effect for VAChT suggests that some components of this profile are stage-dependent rather than stable. Direct *in vivo* PET evidence for cortical opioid and cholinergic receptor systems in schizophrenia is sparse, and no cortex-wide receptor imaging study has examined longitudinal change in these systems, leaving the present findings without a direct empirical comparator. Nevertheless, this gap also points to a genuine contribution of the approach: by extending molecular inference beyond the dopaminergic and glutamatergic systems that have dominated schizophrenia neuroimaging ^23–25^, the FC-informed profiles implicate cholinergic and opioid signaling as candidates for longitudinal molecular change that direct PET measurement has not yet been able to address at the cortical level. If stage-dependent alterations in these systems are confirmed with paired PET-MRI, they would meaningfully expand the neurochemical framework of schizophrenia beyond its classical transmitter hypotheses.

### Limitations and future directions

Several boundaries define the current scope of the framework and motivate specific next steps. First, and most critically for clinical applications, individual-level validation is currently available only for glucose metabolism, amyloid-β, and tau, where paired PET-MRI datasets exist. For the eight neurotransmitter receptor and transporter maps of primary interest for psychiatric research, validation remains at the group-template level: FC eigenmodes reconstruct normative PET atlases beyond spatial autocorrelation, but whether subject-specific differences in reconstructed profiles reflect true individual variation in receptor density is unknown. No dataset currently provides paired individual-level PET and resting-state MRI for these tracers at sufficient sample size, and acquiring such data is a critical next step for establishing the individual-level fidelity of EigenBrain molecular profiles.

Second, the framework relies on a group-level linear model that assumes the relationship between connectome eigenmodes and molecular topography is sufficiently consistent across individuals to transfer learned weights to new subjects. This assumption remains to be formally tested across diverse populations, age ranges, and clinical conditions. At the same time, if eigenmode-molecular correspondence reflects a shared organizational principle of the brain, this consistency would be a conceptual strength rather than a constraint: group-trained weights that generalize across individuals and populations would make the framework scalable and clinically tractable without requiring paired PET-MRI data in every new cohort.

Third, the framework is currently implemented using the Glasser cortical parcellation, which was retained as the primary choice because it integrates multimodal anatomical and functional information to define parcel boundaries, a property not shared by single-modality atlases such as Schaefer or Destrieux. Vertex-level implementations, while appealing in principle for preserving finer-grained spatial detail, introduce non-trivial methodological challenges on both the MRI and molecular sides. On the MRI side, the appropriate number of eigenvector components for convergence at higher spatial resolution is unknown, and whether the elbow criterion established at the parcellation level transfers to vertex-level data requires systematic investigation. On the molecular side, PET-derived receptor maps are inherently limited in spatial resolution by radiotracer binding kinetics, partial volume effects, and point spread function blurring, meaning that at very high vertex resolution, signal from one region contaminates adjacent vertices, introducing spatial contamination that the molecular map itself cannot support. The optimal spatial resolution for eigenmode-based molecular reconstruction therefore remains an open question that constitutes a separate investigation beyond the scope of the present work. In addition, many neuromodulatory systems originate in subcortical nuclei, and disease-related molecular change is often prominent in hippocampal and subcortical regions that the Glasser parcellation excludes; extending the approach to cortico-subcortical systems is therefore a key priority, particularly given that existing PET studies in autism and schizophrenia have been concentrated in subcortical regions.

Fourth, recoverability testing should be performed within each new dataset using sample-specific eigenvectors and the same spatial-null framework, because additional systems may be recoverable in a sample-specific setting beyond the eight validated here with HCP group-level templates.

## Conclusions

The present findings establish that functional, rather than structural or geometric, eigenmodes more faithfully encode cortical molecular topographies, and that this advantage is system-specific: only FC eigenmodes reconstruct a defined subset of neuromodulatory topographies beyond spatial smoothness, while structural and geometric eigenmodes do not. The reconstruction is cross-sample, with molecular weights learned from normative HCP connectomes transferring to independent clinical datasets, providing a biologically validated filter for prioritizing which molecular systems are tractable targets for population-level inference. Looking forward, paired PET-MRI data can be used to learn subject-specific deviations from the EigenBrain baseline, combining eigenmode interpretability with the flexibility of nonlinear prediction, and as clinical MRI scales across multi-site consortia and longitudinal cohorts, EigenBrain offers a translational bridge for molecularly informed stratification and hypothesis generation in populations where direct molecular imaging remains inaccessible.

## Methods

### Datasets and imaging preprocessing

#### Human Connectome Project (HCP)

To generate the group-level multimodal MRI template, we used 94 out of 100 unrelated subjects from the HCP ^53,54^. Six subjects were excluded due to the lack of sessions for rs-fMRI or DWI. MRI data were acquired on the HCP’s custom 3 T Siemens Skyra equipped with a 32-channel head coil. Two T1w images with identical parameters were acquired using a 3D-MP-RAGE sequence (0.7 mm isovoxels, matrix = 320 × 320, 256 sagittal slices; TR = 2,400 ms, TE = 2.14 ms, TI = 1000 ms, FA = 8°; iPAT = 2). DWI was acquired using a single-shot, multiband-accelerated (factor 3) spin-echo EPI sequence (TR = 5,520 ms; TE = 89.5 ms; voxel size = 1.25 mm isotropic), with three diffusion shells (*b* = 1,000, 2,000, and 3,000 s/mm^2^), 90 diffusion directions per shell, and 18 *b* = 0 volumes. Each shell was acquired twice with reversed left-to-right and right-to-left phase-encoding directions. The rs-fMRI was acquired with a single-shot, multiband-accelerated (factor 8) gradient-echo EPI sequence at 2 mm isotropic resolution (TR = 720 ms, TE = 33.1 ms, FA = 52°). Each subject completed two sessions on separate days, each comprising two ∼15-min runs (1,200 volumes per run) acquired with opposite (left–right and right–left) phase encoding, for ∼1 hour of resting-state data in total. The T1w and DWI data were preprocessed with MICApipe v0.2.3 ^70^ to generate GD and SC. For GD, MICApipe identifies a central vertex for each parcel, the vertex with the shortest summed Euclidean distance to all other vertices in that parcel, then computes the GD from this central vertex to every other vertex along the midsurface mesh using Connectome Workbench’s -surface-geodesic-distance, and averages the resulting distances within each parcel. For SC, MICApipe generated a whole-brain tractogram with 10 million streamlines using the iFOD2 algorithm and constructed the structural connectome from cortical parcellations nonlinearly registered to native DWI space. For the FC, we used the HCP minimally preprocessed time series from all four sessions (day1_LR, day1_RL, day2_LR, and day2_RL). For each session, we computed pairwise Pearson correlations between regional time series, applied a Fisher *r*-to-*z* transformation, and then averaged the resulting *z*-matrices across the four sessions to yield a single FC matrix per subject.

#### Neuromaps

The 20 neurotransmitter receptor and transporter maps were from Neuromaps ^34^, previously described by ^8^. The original sources are listed below for the group-level maps.

| Marker | Tracer | *n* | Age (years) |
|---|---|---|---|
| 5-HT<sub>1a</sub> <sup>71</sup> | [<sup>11</sup>C]WAY-100635 | 35 | 26.3 ± 5.2 |
| 5-HT<sub>1b</sub> <sup>72</sup> | [<sup>11</sup>C]P943 | 23 | 28.7 ± 7 |
| 5-HT<sub>2a</sub> <sup>71</sup> | [<sup>18</sup>F]altanserin | 19 | 28.2 ± 5.7 |
| 5-HT<sub>4</sub> <sup>73</sup> | [<sup>11</sup>C]SB207145 | 59 | 25.9 ± 5.3 |
| 5-HT<sub>6</sub> <sup>74</sup> | [<sup>11</sup>C]GSK215083 | 30 | 36.6 ± 9.04 |
| 5-HTT<sup>71</sup> | [<sup>11</sup>C]DASB | 18 | 30.5 ± 9.5 |
| D<sub>1</sub> <sup>75</sup> | [<sup>11</sup>C]SCH23390 | 13 | 33 ± 13 |
| D<sub>2</sub> <sup>76</sup> | [<sup>18</sup>F]fallypride | 49 | 18.41 ± 0.57 |
| DAT<sup>77</sup> | [<sup>123</sup>I]FP-CIT SPECT | 174 | 61 ± 11 |
| NET<sup>78</sup> | [<sup>11</sup>C]methylreboxetine | 10 | 33 |
| H<sub>3</sub> <sup>79</sup> | [<sup>11</sup>C]GSK189254 | 8 | 31.69 ± 8.95 |
| α<sub>4</sub>β<sub>2</sub> <sup>80</sup> | [<sup>18</sup>F]flubatine | 30 | 30.50 ± 10.71 |
| M<sub>1</sub> <sup>81</sup> | [<sup>11</sup>C]LSN3172176 | 24 | 40.45 ± 11.71 |
| VAChT<sup>82</sup> | [<sup>18</sup>F]FEOBV | 18 | 66.8 ± 6.8 |
| CB<sub>1</sub> <sup>83</sup> | [<sup>11</sup>C]OMAR | 77 | 30.01 ± 8.87 |
| MOR<sup>84</sup> | [<sup>11</sup>C]carfentanil | 204 | 32.3 ± 10.8 |
| NMDA<sup>85</sup> | [<sup>18</sup>F]GE-179 | 29 | 40.9 ± 12.7 |
| mGluR<sub>5</sub> <sup>86</sup> | [<sup>11</sup>C]ABP688 | 73 | 19.9 ± 3.04 |
| GABA<sub>A</sub> <sup>87</sup> | [<sup>11</sup>C]flumazenil | 16 | 26.6 ± 8 |
| KOR<sup>88</sup> | [<sup>11</sup>C]LY2795050 | 28 | 33.5 ± 11.3 |

#### Munich Energy Cost

We downloaded Munich datasets (ds004513) from OpenNeuro ^55^. Data were collected at the Technical University of Munich and comprised an experimental cohort of 9 participants (4 females; age, 43 ± 7 years) and a replication cohort of 11 participants (6 females; age, 27 ± 5 years). One participant from the replication cohort was excluded because visual quality control revealed a markedly asymmetric CMRglc map, resulting in a final analytical sample of 19 participants. Imaging was performed using an integrated 3T Siemens Biograph mMR PET/MR scanner (Siemens Healthineers, Erlangen, Germany) with a 12-channel phased-array head coil.

T1w anatomical images were acquired using a three-dimensional magnetization-prepared rapid gradient-echo sequence (3D-MPRAGE; 256 slices; TR: 2,300 ms; TE: 2.98 ms; FA: 9°; FOV: 256 × 240 mm^2^; voxel size: 1 × 1 × 1 mm^3^). DWI were acquired along 30 non-collinear directions at *b* = 800 s/mm^2^, together with one non-diffusion-weighted (*b* = 0) volume (TR: 10,800 ms; TE: 82 ms; voxel size = 2 × 2 × 2 mm^3^). Rs-fMRI was acquired over 10 min using a single-shot echo-planar imaging sequence (300 volumes; 35 slices; TR: 2000 ms; TE: 30 ms; FA: 90°; FOV: 192 × 192 mm^2^; voxel size = 3 × 3 × 3.6 mm^3^).

[^18^F]FDG-PET data were acquired in list mode following intravenous bolus administration of 184 ± 12 MBq of [^18^F]FDG. During PET acquisition, arterial blood radioactivity was measured from the radial artery at 1s intervals using a Twilite blood sampler (Swisstrace, Zurich, Switzerland). Dynamic PET data were reconstructed into 33 frames: 10 × 12 s, 8 × 30 s, 8 × 60 s, 2 × 180 s, and 5 × 300 s. Attenuation correction was performed using T1-derived pseudo-CT images.

Multimodal MRI data were processed using MICApipe following our previously described workflow ^43^. T1-weighted images underwent intensity non-uniformity correction, brain extraction, tissue segmentation, and cortical surface reconstruction. The DWI data were denoised and corrected for Gibbs-ringing artefacts, eddy-current distortions, head motion, and intensity inhomogeneity. The corrected diffusion images were registered to the T1w anatomical image, and tissue response functions and fiber-orientation distributions were estimated using constrained spherical deconvolution. Whole-brain anatomically constrained tractography was then performed, followed by streamline weighting and aggregation within the cortical parcellation to generate participant-specific SC matrices. Rs-fMRI data underwent head-motion correction, nuisance-signal regression, temporal filtering, and registration to the anatomical image. The preprocessed BOLD time series were projected onto the cortical surface and averaged within each cortical parcel, and pairwise temporal correlations were calculated to construct participant-specific FC matrices. Cortical GD matrices were computed from the reconstructed surfaces. SC, FC, and GD matrices were represented using the same cortical parcellation before the EigenBrain analysis. PET images underwent motion correction, 6-mm full-width at half-maximum spatial smoothing, and partial-volume correction using gray matter, white matter, and cerebrospinal fluid masks derived from the T1-weighted images. The CMRglc map was calculated by multiplying the net influx rate constant (*K*_i_) by the plasma glucose concentration and dividing the resulting value by the [^18^F]FDG lumped constant.

#### Dallas Lifespan Brain Study

We downloaded Dallas Lifespan Brain Study (ds004856) from OpenNeuro ^56^. The dataset comprises three longitudinal assessment waves. The present study included 61 participants from wave 3 with contemporaneous T1w, DWI, rs-fMRI, tau PET, and amyloid-β PET data. The sample included 35 females and 26 males, with ages ranging from 39 to 96 years (mean ± s.d., 69.3 ± 13.1 years). Cognitive impairment was assessed using the Alzheimer’s Disease Assessment Scale (ADAS). ADAS total scores ranged from 1 to 12 (mean ± s.d., 5.5 ± 2.8).

T1w images were acquired using an MPRAGE sequence (160 sagittal slices; TR: 8.1 ms; TE: 3.7 ms; FA: 12°; voxel size: 1 × 1 × 1 mm^3^). Rs-fMRI used a T2*-weighted echo-planar imaging sequence (43 slices; TR: 2,000 ms; TE: 25 ms; FA: 80°; voxel size: 3.4 × 3.4 × 3.5 mm^3^). DWI was acquired along 30 non-collinear directions at *b* = 1,000 s/mm^2^, with one *b* = 0 volume (TR: 4,410 ms; TE: 51 ms; voxel size = 1.75 × 1.75 × 2 mm^3^). Amyloid-β PET was acquired using [^18^F]florbetapir ([^18^F]AV-45; approximately 370 MBq), with two 5-min emission frames beginning 50 min after injection. Tau PET was acquired using [^18^F]flortaucipir ([^18^F]AV-1451; approximately 370 MBq), with a 20-min scan beginning 80 min after injection. PET imaging at wave 3 was performed using a Siemens Biograph mCT Flow Edge scanner.

Multimodal MRI data were processed using MICApipe (using the preprocessing described above) to derive cortical GD, SC, and FC matrices. Amyloid-β and tau PET were also processed as described above. Amyloid-β and tau deposition were quantified using standardized uptake value ratio (SUVR) maps with whole-cerebellum reference.

#### Autism

We included 140 autistic individuals and 143 non-autistic controls (all male) from five ABIDE cohorts (NYU-I, NYU-II, PITT, USM, and TCD), ranging in age from 5.3 to 39.4 years (mean ± s.d., 17.6 ± 7.4 years). Site-specific acquisition parameters are available through the ABIDE data portal https://fcon_1000.projects.nitrc.org/indi/abide/. Full preprocessing details are provided in ^61^. Briefly, T1-weighted images were processed using FreeSurfer v5.1. Resting-state fMRI data were obtained from the Preprocessed Connectomes Project and processed using C-PAC, including slice-timing correction, motion correction, CompCor-based nuisance regression, band-pass filtering (0.01-0.10 Hz), and registration to MNI152 space.

#### Geneva-Berlin

The longitudinal dataset was collected at two sites, Geneva and Berlin. We selected participants who completed MRI and fMRI scanning at both time points, yielding 42 participants in Geneva (20 individuals with schizophrenia and 22 healthy controls) and 50 in Berlin (27 individuals with schizophrenia and 23 healthy controls). Participants were asked to return for a second scan approximately three months later, with a maximum delay of one week. Of the 92 individuals, 74 were male and 18 were female. Age ranged from 21 to 58 years (mean ± s.d., 38.1 ± 9.0).

Imaging was performed at Berlin (Charité Campus Mitte, Psychiatry CCM; Siemens Prisma-fit) and Geneva (Fondation Campus Biotech Geneva; Siemens Prisma), using 3T scanners running syngo MR E11 with a 64-channel head-neck coil. Both sites used identical acquisition protocols. T1w images were acquired using an MPRAGE sequence (TR: 2,200 ms; TE: 2.96 ms; TI: 900 ms; FA: 9°; voxel size: 1 × 1 × 1 mm^3^; GRAPPA acceleration factor: 2). Rs-fMRI used a T2*-weighted echo-planar imaging sequence (66 axial slices; TR: 1,000 ms; TE: 32 ms; FA: 50°; voxel size: 2 × 2 × 2 mm^3^; multiband acceleration factor: 6). MRI data were processed using MICApipe (using the preprocessing described above) to derive GD and FC matrices.

### Manifold learning

To reduce computational dimensionality and ensure correspondence across modalities, all preprocessed volumetric maps were coregistered to the corresponding T1-weighted images, projected onto the cortical mid-thickness surface, and resampled to fsLR-32k surface space. Surface data were then summarized using the 360-region Glasser cortical parcellation ^57^.

Cortical geodesic distance (GD), structural connectivity (SC), and functional connectivity (FC) were consequently represented as 360 × 360 region-wise matrices.

We applied diffusion map embedding, as described by ^59^ and implemented in BrainSpace ^41^, to obtain low-dimensional representations of each matrix. Briefly, diffusion map embedding converts each matrix into an affinity matrix, constructs a diffusion operator describing relationships among cortical regions, and decomposes this operator into eigenvalues and eigenvectors ^58,89^. Diffusion map embedding is governed by the parameter α, which controls the influence of sampling density on the estimated manifold (α = 0, maximal influence; α = 1, no influence). Following previous recommendations ^41,59,61,90^, we set α = 0.5 to preserve global relationships among cortical regions while reducing sensitivity to noise in the covariance matrix. The diffusion map embedding algorithm is robust to noise and computationally efficient compared to other nonlinear manifold learners ^91,92^. Before embedding, matrices were proportionally thresholded to retain the strongest 10% of connections, consistent with previous studies ^59,61,62,93^. The resulting eigenvectors represent spatial gradients, ordered according to the proportion of spatial variance captured. Slowly varying eigenvectors describe broad cortical organizational axes, whereas higher-order eigenvectors capture progressively finer spatial patterns. Based on previous evidence that approximately 70 components are sufficient for map reconstruction to converge ^43^, we retained the first 100 non-trivial eigenvectors from each modality to ensure adequate model coverage. These eigenvectors served as spatial basis functions for molecular-map reconstruction. Because eigenvector orientation is mathematically arbitrary, individual-level eigenvectors were aligned to the corresponding group-level HCP reference using Procrustes rotation before subsequent analyses.

### Molecular map reconstruction

EigenBrain reconstructs a molecular map as a weighted linear combination of MRI-derived connectomic eigenvectors. For each molecular marker, regional values across the 360 Glasser parcels formed the response vector, and the corresponding GD, SC, or FC eigenvectors formed the predictor matrix. Model weights were estimated using ordinary least-squares (OLS) regression:

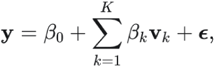

where ***y*** is the observed molecular map, ***v****_k_* is the *k*-th eigenvector, *K* is the number of included components, and *β_k_* is its fitted weight. The reconstructed map was obtained by applying the estimated weights to the corresponding eigenvectors.

We fitted three single-modality models based on GD, SC, or FC and three multimodal models: GD_base_ + FC, SC_base_ + FC, and GD_base_ + SC_base_ + FC. For each single-modality model, components were entered hierarchically according to their eigenvalue rank, from 1 to 100. Reconstruction performance was evaluated at each model order using the coefficient of determination (R^2^) and the Pearson correlation between observed and reconstructed molecular maps. The “base” for GD or SC was defined as the number of components at the elbow of its component-number/R^2^ curve, representing the point at which additional components produced only limited improvement. In multimodal models, these GD and SC bases were held fixed while FC components were added hierarchically.

Group-level models for the 20 neurotransmitter receptor and transporter maps were trained using molecular templates from Neuromaps and eigenvectors derived from the unrelated HCP sample. For individual-level reconstruction, participant-specific eigenvectors were aligned to the corresponding HCP reference using Procrustes rotation. The group-level model coefficients were then transferred to the aligned individual eigenvectors to generate participant-specific molecular profiles.

For glucose metabolism, amyloid-β, and tau, models were trained using the group-mean PET maps and HCP-derived eigenvectors. Individual reconstructions were generated using either HCP-derived GD and SC combined with participant-specific FC or participant-specific GD, SC, and FC. Individual-level accuracy was quantified using the Pearson correlation between reconstructed and observed PET maps across cortical parcels.

### Statistical analysis

All statistical analyses were performed using Statsmodels (https://www.statsmodels.org/). In the autism sample, parcel-wise case-control differences were assessed using OLS:

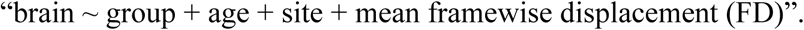

In the longitudinal schizophrenia cohort, generalized estimating equations (GEE) were used to account for repeated measurements within participants. Parcel-wise longitudinal effects were tested using:

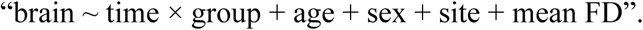

Spatial correspondence between brain maps was quantified using Pearson correlation. Statistical significance was assessed against 1,000 surrogate maps generated using spatial variogram models that preserve the spatial autocorrelation of the empirical maps ^41,60^. Surrogates were generated using the Glasser atlas GD matrix, and the resulting spatially corrected significance values are reported as *P*_variogram_. In addition, to test whether reconstruction accuracy exceeded that expected from spatial autocorrelation alone, we generated 1,000 surrogate maps for each molecular marker using spatial variogram models that preserve the empirical spatial autocorrelation and compared the reconstruction accuracy using 20, 40, 60, 80, and 100 eigenvectors\ of each true map against the distribution obtained across its surrogates. This analysis was run for the single-modality GD, SC, and FC models using HCP group-level eigenvectors.

## Data availability

All publicly available datasets used in this study are described in the Datasets section and can be downloaded from their respective project websites or OpenNeuro. The longitudinal Geneva-Berlin dataset was collected locally and is available from Stefan Kaiser and Florian Schlagenhauf upon reasonable request, subject to applicable ethical and data-protection requirements.

## Code availability

Data preprocessing and statistical scripts for this study are shared at a GitHub repository: https://github.com/wanb-psych/EigenBrain_framework.

## Competing Interests

Dr. Richard A.I. Bethlehem is a founder of and holds equity in Centile Bioscience Inc. Dr. Matthias Kirschner has received consulting fees from Otsuka for activities unrelated to the present study. All other authors declare no competing interests.

## Acknowledgements

Bin Wan acknowledges funding from the Novartis Foundation for Medical-Biological Research (No. 26B040). Matthias Kirschner acknowledges funding from the Swiss National Science Foundation (No. 32003B_219240). Sofie Valk gratefully acknowledges support by the “Precision Medicine for Mental Health” research focus of University Medicine Zurich (UMZH); ERC starting grant (No. 101220063); Hector Research Foundation; Jacobs Research Foundation.

## Authors’ contributions

**Conceptualization:** B.W.; **Methodology:** B.W.; **Formal analysis:** B.W.; **Writing - Original Draft:** B.W., M.Kirschner; **Writing - Review & Editing:** All authors; **Visualization:** B.W.; **Data curation:** B.W., J.Y.H., Z.Q.L., S.J.H, F.C., B.M., F.S., S.K., M.Kirschner; **Project administration:** B.W., **Funding acquisition:** B.W., M.Kirschner; **Supervision:** B.W., M.Kirschner.

